# Extracellular matrix guidance determines proteolytic and non-proteolytic cancer cell patterning

**DOI:** 10.1101/2022.03.16.484647

**Authors:** Lianne Beunk, Sjoerd van Helvert, Bram Bekker, Lars Ran, Ross Kang, Tom Paulat, Simon Syga, Andreas Deutsch, Peter Friedl, Katarina Wolf

**Author notes:** Corresponding author: Dr. Katarina Wolf Dept. of Cell Biology, Radboud Institute for Molecular Life Sciences (RIMLS), Geert Grooteplein Zuid 26-28, 6525 GA Nijmegen.

## Abstract

Metastatic tumor cell invasion into interstitial tissue is a mechanochemical process that responds to tissue cues and further involves proteolytic remodeling of the tumor stroma. How matrix density, tissue guidance and the ability of proteolytic tissue remodeling cooperate and determine decision-making of invading tumor cells in complex-structured three-dimensional (3D) tissue remains unclear. We here developed a collagen-based invasion assay containing a guiding interface of low collagen density adjacent to randomly organized 3D fibrillar lattice and examined the invasion of melanoma cells from multicellular spheroids in response to matrix density, guidance cues and collagenolysis. After 48 hours of culture, two invasion niches developed, (i) sheet-like collective migration along the interface and (ii) single cell- and strand-like invasion into randomly organized 3D matrix. High collagen density impeded migration into the random matrix, whereas migration along a high-density collagen interface was increased compared to the low-density matrix assay. *In silico* analysis predicted that facilitated interface migration in high-density matrix depended on physical guidance without collagen degradation, whereas migration into randomly organized matrix was strongly dependent on collagenolysis. When tested in 3D culture, inhibition of matrix metalloprotease (MMP)-mediated collagen degradation compromised migration into random matrix in dependence of density, whereas interface-guided migration remained effective. In conclusion, with increasing tissue density, matrix cues bordered by dense matrix, but not randomly organized matrix, support effective MMP-independent migration. This identifies the topology of interstitial tissue a primary determinant of switch behaviors between MMP-dependent and MMP-independent cancer cell invasion.

## Introduction

Interstitial tissues connect and support organ-specific tissues in the body and are rich of extracellular matrix components, with collagen forming predominant supportive architectures. Due to their variety in composition and function, interstitial compartments are highly heterogenous in density, topology and stiffness [1,2]. In the interstitium of the upper skin for example, collagen fibers form structures that span one dimensional (1D) line-like single-fiber structures as well as three dimensional (3D) random network-like topologies in addition to components such as blood vessels, nerves or glands [3–5]. In contrast, the lower dermis consists of dense collagen of interwoven stiffer cable-like structures, and additional tissues like muscular layers or perineural tracks offer long linear structures [4,5]. These tissue architectures form topologies that contain narrow cleft-like interfaces and tunnels of linear geometry of often hundreds or thousands of micrometer length that are filled with interstitial fluid and with cross-sectional diameters ranging between 0.5 and 30 μm [4–6]. Overall, interstitial tissues form heterogeneous structures that, in the context of cell migration, contain both barriers as well as linear guidance cues. The complex extracellular matrix (ECM) architecture can therefore function as both an inhibitor or promotor of cell locomotion.

Cell migration is essential for many physiological processes including immune response, tissue remodeling and wound healing, but also contributes to pathology such as during cancer invasion and metastasis, including melanoma progression [7–9]. During migration within dense porous substrate, mesenchymal cells need to negotiate for space, either by adapting their shape including the spacious and stiff nucleus to pre-existing space or by proteolytic ECM removal [10,11]. Degradation of interstitial collagen is mediated mainly by matrix metalloproteinases (MMPs), with their expression being upregulated upon progression in many tumors including melanoma [12,13]. Many tumor cell types, including melanoma cells, employ pericellular proteolysis during migration to overcome resistance imposed by high-density matrix, hereby remodeling the extracellular matrix [14–16]. As a result, barrier-free tracks are generated which are filled by follower cells during proteolytic tumor cell invasion into dense collagen networks [15,17]. This process results in the formation of collective cell strands surrounded by dense ECM that, depending on the cells origin, may form functional cell-cell junctions, such as mediated by E-cadherins, ALCAM or other candidate adhesion systems [14,15,18,19]. Consistently, tumor-derived collagenolytic remodeling assists in partly overcoming matrix resistance and enhancing cell invasion into dense surrounding tissue that laterally confines and guides the cell [14]. When proteolysis is low or absent, in particular in strongly confining substrates, cell motility is abrogated [14,19]. Thus, dense 3D ECM forms a barrier towards migrating cells that can be overcome by proteolysis and become remodeled into a guiding ECM structure.

Track-like tunnels and clefts *in vivo* and *in vitro* provide linear cues to migrating cells that serve as guidance cues and enhance cell motility efficacy [19–23]. Depending on the density and stiffness, and thus compressibility, of the surrounding tissues, spaces may widen to some extent and be able to harbor migrating cells. Guided by tissue tracks, migrating cells, even in the absence of proteolysis, may fill and physically widen pre-existing barrier-free, yet confining, tracks in a collective pattern, consequently forming a collective migration phenotype independent on proteolysis [19,24,25]. Whereas cell patterning into either tissue compartment was studied before, how tumor cells chose ECM patterns for 3D invasion depending on guidance cues and the ability to proteolytically degrade it, is yet largely unknown.

To dissect whether tumor cells can choose between MMP-dependent or MMP-independent migration routes, we here developed a collagen-based *in vitro* assay that contained both random collagen networks at different collagen concentrations, as well as a low-density guiding interface. Using a MV3 melanoma spheroid model, we monitored the relevance of tumor cell mediated collagenolysis on cell migration in dependence of ECM geometry and density. Our results indicate that pronounced linear cues within high ECM density enable non-proteolytic migration in an otherwise migration-prohibitive environment.

## Methods

### Antibodies and inhibitors

Affinity purified rabbit anti-collagen type I cleavage site antibody (Col1¾C_short_, Immunoglobe), Alexa-488-conjugated secondary pre-absorbed goat anti-rabbit IgG (Invitrogen), DAPI (Sigma) and Alexa-568-conjugated phalloidin (Invitrogen) were used. To block collagen degradation, the broad-spectrum MMP inhibitor GM6001 (dissolved in DMSO; Calbiochem) at an end concentration of 5 μM was used.

### Cell culture and spheroid generation

Human wild-type MV3 melanoma cells (kindly provided by G. van Muijen, Dept. of Pathology, Radboudumc Nijmegen) were cultured in Dulbecco’s modified Eagle’s serum medium (DMEM, Invitrogen) supplemented with 10% fetal calf serum (FCS, Sigma), penicillin (100 U/ml, PAA), streptomycin (100 μg/ml, Invitrogen), L-glutamine (2 mM, Lonza) and sodium pyruvate (1 mM, Gibco) and maintained in a humidified 10% CO_2_ incubator at 37°C. Cells were detached with EDTA (2 mM, Invitrogen) and multicellular spheroids were generated according to the hanging drop method [26]. Briefly, hanging droplets of a 30 μl volume containing 5000 cells, methylcellulose (40% dissolved in DMEM; Sigma) and bovine collagen (10 μg/mL; Advanced Biomatrix) were incubated for 24 hours to ensure multicellular aggregation. Spheroids were washed and resuspended in medium.

### 3D spheroid collagen-collagen interface culture

To prepare a spheroid-collagen culture, square silicon molds (Electron Microscopy Sciences) were attached to the bottom of a 24 well culture plate. Fibrillar ‘high-density’ (6 mg/ml) or ‘low-density’(2 mg/ml) collagen gels were constituted by using rat tail collagen type I (Corning) supplemented with 10x Minimum Essential Serum (Sigma) and neutralized to a pH of 7.4 with 1N NaOH, pipetted into the mold, and polymerized at 37°C. Where indicated, a single spheroid was added on top of the first gel after polymerization, the excess medium removed, and a second collagen droplet added on top and polymerized, and covered with medium. Where indicated, GM6001 or DMSO were supplemented to both collagen lattices and supernatant. Spheroid-containing 3D interface lattices were imaged within 1 hour after collagen polymerization and further incubated for 48 hours, fixed with 4% PB-buffered PFA, washed and stained with indicated antibodies and fluorescent dyes, and XY imaging was performed. Afterwards, gels were manually cross-sectioned using a scalpel during concomitant inspection under a brightfield microscope. Cross-sectioning occurred from one side of the spheroid and was repeated until the spheroid core was cut in the middle and the resulting lattice containing half the spheroid, the remaining lattice was then turned by 90° and imaged resulting in an XZ image depiction of the spheroid. Where indicated, interface-containing collagen lattices without cells were fixed within 1 hour after polymerization, washed, stained and imaged in XY or XZ direction.

### Microscopy

Imaging was generally performed using high-content brightfield and epifluorescence microscopy combined with automated multi-position image acquisition and stitching (Leica DMI6000B, 10x / 0.25 NA air objective, 10 μm z-interval, 24 slices). For higher resolution detection of collagen, cells and collagen cleavage epitope, confocal microscopy (Zeiss LSM880, 10x / 0.45 NA air objective; Olympus FV1000, 40x / 0.8 NA water objective) was used. Sor high resolution detection of the collagen interface by scanning electron microscopy (SEM), cell-free samples were fixed for 1 hour at room temperature with 2% glutaraldehyde (Merck) in 0.1M cacodylate buffer, followed by two washes in the same buffer. 1% osmium tetroxide (EMS) in 0.1M cacodylate buffer was used as a secondary fixative with incubation for 1 hour at room temperature. Samples were manually cross-sectioned, then processed for SEM by dehydration through a graded series of ethanol followed by critical point drying using CO_2_ and then sputtered with gold and imaged by SEM (JSM-6340F; JEOL).

### Image analysis

Image processing and quantification was performed by Fiji ImageJ (1.52n; National Institutes of Health). Images were cropped, rotated, manually adjusted for contrast and brightness, and displayed in virtual colors, from which readouts for guided migration (migration area, cell number, maximally migrated distance) and non-guided 3D migration (migrated distance and cell numbers) were performed. The migration area was quantified using brightfield images taken within 1 hour after polymerization as well after 48h. The images were processed by automated thresholding of the cell-populated area using the Triangle method, followed by filling of artificial holes, quantifying the areas at 0 hour and 48 hours excluding areas smaller than 100 μm^2^, and afterwards forming the difference (Figs. S2A, B). Cell numbers in the migration area were quantified in a single slice, capturing all cells in the interface. After the spheroid center was manually excluded, particle analysis by using a median filter, automated thresholding (Default method) and watershedding of the DAPI signal was performed and areas larger than 1000 μm^2^ were excluded (Fig. S2D). The maximum migration distance was quantified by measuring the distance from the rim of the core to the 6 furthest leader cells and averaged per spheroid (Fig. S2E). To quantify non-guided migration into the random collagen lattices, maximum intensity projections of the DAPI channel after manual cross-sectioning were used. The number of cells present within each zone, determined by a ROI that was expanded by each 25 μm around the rim of the spheroid, were counted manually using the Fiji Cell Counter plugin (Figs. 1L; S3B). The imaged cross-sectioned lattice only contained half of the spheroid and the imaging penetration depth was smaller than the non-guided migration distance that was measured. Taken together, not all cells within the 3D matrix were imaged, using the penetration depth, non-guided migration depth and core size, it was calculated that only 40.4% of non-guided cells were imaged. Consequently, the total number of non-guided cells was corrected with a factor 2.47 (Figs. S3A,C,D). Further, to quantify the reflection signal of the interface as compared to the 3D collagen lattice, mean gray values from lines of 1500 pixel width were normalized to the intensities in random 3D matrix, and from different samples the lowest signal at the interfaces were overlaid (Fig. S1). Last, the COL1¾ signal was calculated as mean gray signal from cell-populated areas and divided by the number of cell nuclei after subtraction of the background signal (Fig. 4A).

**Figure 1.**
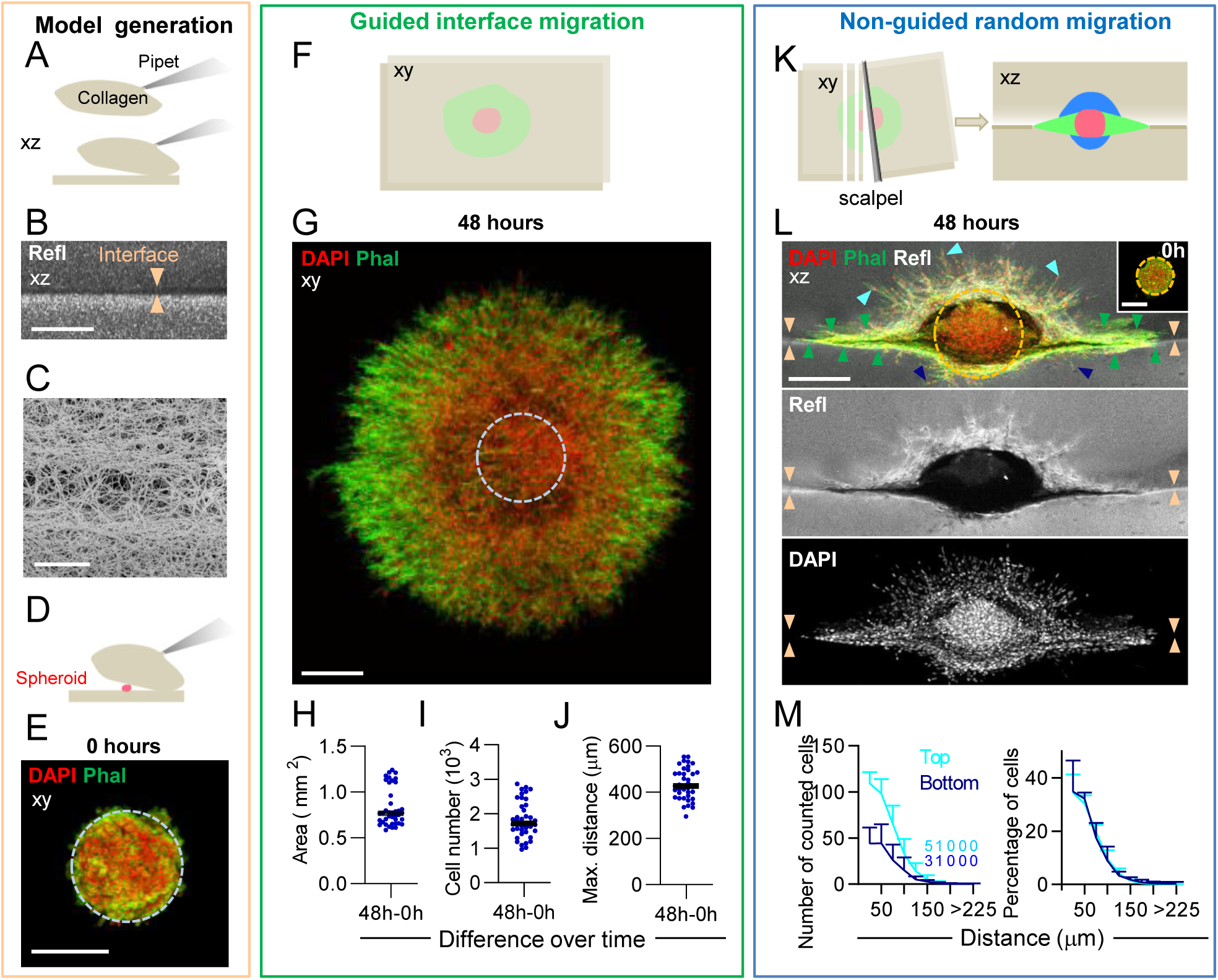
Emigration of MV3 melanoma cells from spheroids into the high density 3D interface model. The collagen-collagen interface model **(A-C)**, with embedded spheroids after 0h **(D**,**E)** or 48h **(F-M)** culture. **(A)** Cartoon depicting model generation at the beginning of the experiment. **(B**,**C)** Depiction of the collagen-collagen interface (arrowheads) consisting of low collagen density between two high-density (each 6 mg/ml) collagen lattices by confocal reflection microscopy **(B)** or SEM **(C)**. xz-side view. **(D)** Cartoon depicting spheroid embedding. **(E)** Spheroid morphology directly after collagen polymerization. Cells and collagen were visualized as indicated (DAPI; Phal, phalloidin) and are shown as top view (xy) Dotted circle indicates average spheroid size at 0h. **(F)** Schematics of emigration into the interface (green) seen from the top. **(G)** Cell emigration into the interface. Dotted circle, outline of the original spheroid core (see D). **(H-J)** Quantification of area, cell number and maximally migrated distance of cells outside the spheroid core (see Fig. S2 for analysis procedure). Data represent 2-10 measurements (dots) per experiment (N=6). **(K)** Schematics depicting the manual cross-section method and resulting side view of interface-guided cell emigration pattern (green) and migration into the porous random collagen network (blue); spheroid core (red). **(L)** Cell emigration from spheroid into the interface (green arrowheads), and into the lower (dark blue arrowheads) and upper (light blue arrowheads) random collagen lattices. Orange arrowheads indicate the interface, dotted circle, average initial spheroid size (see upper inset for spheroid size at 0h culture). **(M)** Quantification of cells located in the upper and lower random collagen lattices as a function of migrated distance from the spheroid core (see Fig. S3). Left, absolute number of counted cells; numbers within the graph depict the number of cells that were located at the indicated distance from the spheroid. Right, distribution of cells into the lower and upper collagen compartment. Data represent 2-10 spheroids per experiment (N=6). **(H-J)** Solid line, median, **(M)** solid line, mean; whiskers, SD. Scale bars: 250 μm, except **(C)**: 50 μm.

### *In silico* model

To predict the effects of collagen density and of collagenolysis on cell migration, we developed an *in silico* cellular automaton. The detailed mathematical procedures, simulation code and simulation videos were deposited at GitHub (https://github.com/TomPlt/Collagen-Interface-Model). Cell density and ECM density were represented by scalar fields defined on a 3D lattice with a lattice spacing of 10 μm (Fig. 3A(I)). Cell migration into the interface and the random matrix was determined by a deterministic process depending on the local, spatial microenvironment defined as the six neighboring lattice sites (Von Neumann neighborhood; Fig. 3A(II)). We initiated the lattice with a spherical volume of high cell density representing the spheroid and a plane with low ECM density surrounded by high ECM density as an interface, mimicking the experimental setup. The system was simulated for 100 discrete time steps in 30 minute intervals closely matching the duration of the experiments (48 hours; Fig. 3B). We assumed that cells were able to degrade surrounding ECM. This was implemented by a linear dependence of ECM degradation on cell density in the voxel and the neighboring voxels. For migration, two basic assumptions were made: (1) Cell migration is limited by available space, which was implemented by a linearly decreasing relation between outward migration rate and cell density in the neighboring voxels. Cells are also hindered to invade high-density ECM regions. (2) Surrounding ECM serves as a substrate for cells allowing them to adjust their movement direction along high-density ECM regions (contact guidance). This complex relation between cell migration and the ECM neighborhood was implemented using a function that increases linearly with ECM density in the neighborhood but decreases exponentially with the ECM density in the target voxel for the case of a homogeneous ECM environment (Fig. 3A(III)). To replicate the experimentally observed relative and absolute cell numbers in the interface for the high- and low-density collagen, we performed broad parameter scans varying through all possible combinations of critical density factor, random ECM density, interface density and degradation constant. The results for the ECM interface densities for the low-density collagen ranged between 0.2 and 0.32, whereas the values for the high-density collagen were significantly lower (Table 1). To simplify following simulations, we set the ECM densities in the interface for both collagens to the mean of the corresponding range (0.26 and 0.1). Repeating the parameter scan with fixed interface densities we obtained the model parameters which were used in all further simulations for both collagen gel densities (Table 1).

**Table 1:** Model parameter values after fitting the model for the experimentally observed migration numbers in the collagen lattices of both indicated concentrations

| Collagen concentration | Interface density | Random 3D ECM density | Degradation constant | Critical density |
| --- | --- | --- | --- | --- |
| **2 mg/ml** | [0.2, 0.32] | [0.46, 0.52] | [0.04, 0.08] | 0.1 |
| **6 mg/ml** | [0.08, 0.12] | [0.74, 1.00] | [0.12, 0.18] | 0.1 |

### Statistics

Statistical analysis was performed by the two-tailed non-paired Mann-Whitney test for samples with non-Gaussian distribution and, when necessary, correction for multiple testing was performed by the Holm-Sidak method using GraphPad Prism 8 software.

## Results

### Induction of dominant cell emigration into the guiding extracellular matrix interface

To obtain a collagen-based substrate that consists of both a porous and a guiding component, we developed an *in vitro* 3D collagen ‘interface’ assay. By polymerizing a high-density 6 mg/ml 3D collagen lattice and subsequently reconstituting a second collagen gel on top (Fig. 1A), a layer of reduced density formed between, but physically connected, the bottom and top lattices (Figs. 1B; S1A,B). When compared to the two neighboring lattices consisting of randomly organized collagen fibrils, the linking interface consisted of a fiber network of around 45% reduced signal intensity (Fig. S1C) and increased porosity (Fig. 1C). In addition, the interface was further supported by a 10-20 μm thin edge in the bottom gel consisting of increased collagen density as indicated by a 3-4 times increased reflection signal (Figs. 1B,C; S1A,C). The mechanism for this asymmetry in density is not known, however differential polymerization kinetics, water evaporation, and effects of gravity may have contributed to condensation of the lower collagen interface. MV3 melanoma spheroids were embedded between the two high-density gel layers (Fig. 1D) and remained intact without detached cells being present directly after gel reconstitution, confirming that spheroid morphology was largely unaffected by the constitution process (Fig. 1E). Thus, a 3D collagen matrix environment comprising a low-density interface adjacent to a randomly polymerized 3D fibrillar network for spheroid culture was developed.

The interface assay enabled us to investigate the effect of extracellular matrix cues on cell patterning over time (Fig. 1F). Melanoma emigration into the interface occurred in a homogeneous sheet-like fashion (Fig. 1G). The area populated by the spheroid after 48 hours of culture was significantly larger as compared to the beginning of the experiment, while the size of the core did not increase (Figs. S2A-C). We therefore here define the cells present in a newly covered area of around 0.75 mm^2^ as cells that have actively emigrated from the spheroid (Figs. 1H; S2A,B). Accordingly, around 2000 cells located outside the core of the spheroid (Figs. 1I; S2D) with a maximally migrated distance of in average 400 μm between the rim of the core and the outer spheroid rim (Figs. 1J; S2E).

To dissect migration efficacies into the different layers of the collagen assay, we established a methodology to cross-section the lattices and image these at a 90° angle (Figs. 1K; S3A). The highly efficient sheet-like migration was located in the cleft-like part of the interface, while emigration into the random matrix both above and below the spheroid center was present in collective strand-like patterns, though at lesser distance (Fig. 1L). Cell migration into the top layer was favored compared to the lower collagen compartment, while we observed a similar decrease in cell patterning from the rim of the spheroid core in a non-linear, non-logarithmic manner for both layers (Figs. 1M; S3B). Presumably, the presence of the high-density collagen layer between the spheroid core and the lower random lattice reduced the number of cells entering into the matrix, but once access was gained, similar migration kinetics occurred (Fig. 1M, right). After application of an imaging-related correction factor (see Image analysis paragraph in the Methods section), 350 and 150 cells migrated into the top and bottom layer, respectively, thus on average 500 cells in total (Figs. S3C,D).

Taken all cells together that left the tumor spheroid, the interface, as compared to the 3D collagen lattices, harbored approximately 3 times more cells (1700 versus 500 cells; Figs. 1I; S3D) that migrated around 2.5 times further (400 versus 150-175 μm; Figs. 1J,M). Thus, compared to the surrounding random matrix, the cleft-like interface in a high-density matrix environment supported prominent migration both in terms of numbers and distance.

### Invasion efficiency in response to ECM density and interface guidance

To dissect how collagen density and guidance patterns impact decision making of invading tumor cells, we examined migration in the high-density (6 mg/ml) interface model. In addition, as a reference for constitutive collagenase-independent migration, an interface-containing low-density (2 mg/ml) collagen lattice was used. Whereas at high collagen density, a clearly defined interface with great differences in collagen density is observed (Figs. S1A-C), decreasing collagen concentration resulted in a less pronounced linear interface structure with less differences in collagen density in the interface and the surrounding random matrix (Figs. S1D-F). While emigration along the cleft occurred as a continuous sheet without defined strand-like substructures in the high-density assay (Fig. 2A), invasion in the interface bordered by low-density matrix resulted in multi-cell strands that in part contained long protrusions at the tip (Fig. 2A, arrowheads). Cell emigration into the low-density interface was reduced compared to the high-density formed interface with regard to area, cell number and maximum locomoted distance (Figs. 2B-D). At low collagen density, effective, predominantly single-cell infiltration into the random, non-guiding compartment was observed (Fig. 2E), which was reflected by an increased migration distance and a higher cell number in the low-density lattice (Figs. 2F,G). Migration of cells along the well-defined interface bordered by high-density ECM is favored as the relative fraction of cells migrating along the interface was decreased in the low-density assay (Fig. 2H).

**Figure 2.**
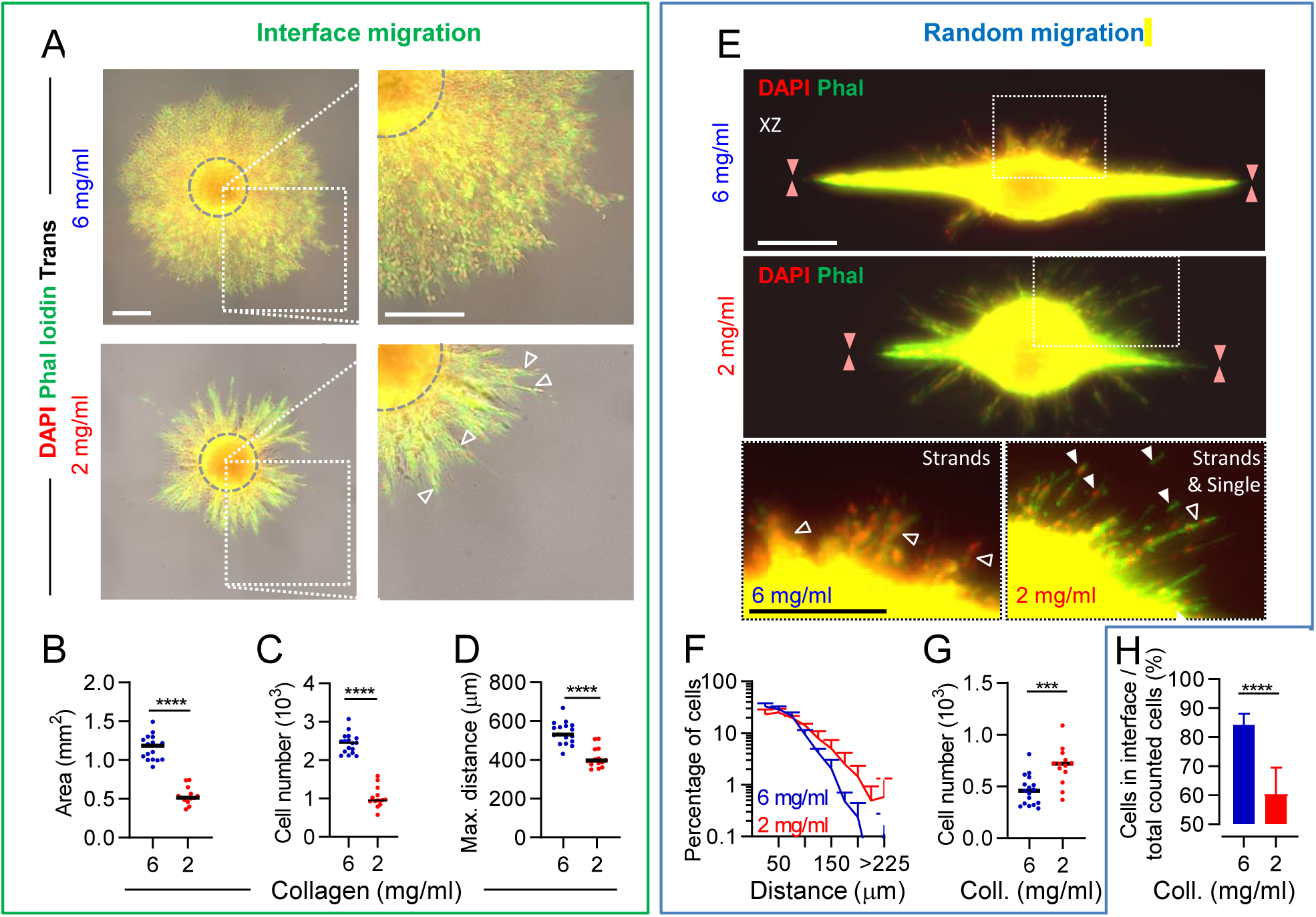
Graded migration in collagen-collagen interface lattices of different densities. Invasion from the spheroid in the interface assay at indicated collagen densities after 48h culture. **(A)** Xy view. Gray dotted circles indicate size of depicted spheroid when imaged at 0h. Trans, transmission. Open arrowheads indicate collective strands. Rectangles indicate the zones for the zoom-ins at the right column. **(B-D)** Quantification of interface-guided migration parameters as indicated (see Fig. 1G-I). Solid line, median. Data represent 1-5 measurements (dots) per condition per experiment (N=4). **(E)** Xz view. Orange arrowheads indicate location of the interface. Open white arrowheads indicate collective strands, and filled white arrowheads indicate single cells. **(F**,**G)** Quantification of **(F)** distribution and **(G)** total numbers of non-guided cells in lattices of indicated concentrations. Data represent 1-5 measurements (line/dots) per condition per experiment (N=4). **(H)** Percentage of cells in the interface to total number (into interface and random collagen) of migrated cells per spheroid. Data represent 1-5 measurements (bar) per condition per experiment (N=4). **(B-D, G)** Solid line, median; **(F**,**H)** solid line or bar, mean; whiskers, SD. Mann-Whitney test, ***: P-value < 0.001; ****: P-value < 0.0001. N=4; 1-5 spheroids per condition per experiment. Scale bars: 250 μm.

### *In silico* analysis of cell invasion in response to ECM density and topology

To identify the role of ECM density along patterned ECM as minimal requirement for guidance within a high-density matrix, we applied an *in silico* cellular automaton model in which cells were represented by scalar cell density values on a 3D lattice (Fig. 3A(I)). The cell densities changed according to a deterministic interaction rule that incorporated proliferation, matrix degradation and guidance by ECM (Figs. 3A(II,III)). We simulated a spheroid moving into a planar linear cleft of low collagen density mimicking the interface between two 3D collagen meshworks (Fig. 3A(I)). Like in the *in vitro* experiments, the interfaces allowed for migration further away from the spheroid than the 3D compartment, resulting in the characteristic spheroid shape also observed *in vitro* (Fig. 3B). As determined by a broad parameter scan (see Methods section), we identified that the high 3D collagen density was accompanied by reduced collagen density between the layers (ECM_int_ = 0.1), whereas the low-density collagen assay results in a less well-defined boundary along the interface with higher interface collagen density (ECM_int_ = 0.26). Thus, a well-defined interface for the high-density condition and a more shallow, less precise interface for the low-density condition was obtained, consistent with the differences in collagen density between the high-density and low-density assay as detected by confocal reflection (Fig. S1).

To examine the effects of random ECM density and degradation rate on the ratio of migration along the guiding interface, the interface density was kept constant, while the density of random ECM and the collagen degradation were varied. The percentage of cells migrating along the interface of all emigrating cells increased with increasing random ECM densities and decreasing degradation constant values (Fig. 3C). Furthermore, for the model to match with the *in vitro* results and to replicate absolute and relative cell numbers in the interface (Figs. 2C,H), cells had to degrade 6 mg/ml collagen around 2.5 times faster compared to 2 mg/ml collagen, which is in good agreement to previous experimental findings [27]. Taken together, the cellular automata model allowed us to predict that cell migration along the interface was favored with increasing surrounding matrix density and decreasing proteolytic activity. In particular, the model predicted that without collagenolysis, migration along the interface between high-density collagen would be most efficient (Fig. 3C).

**Figure 3:**
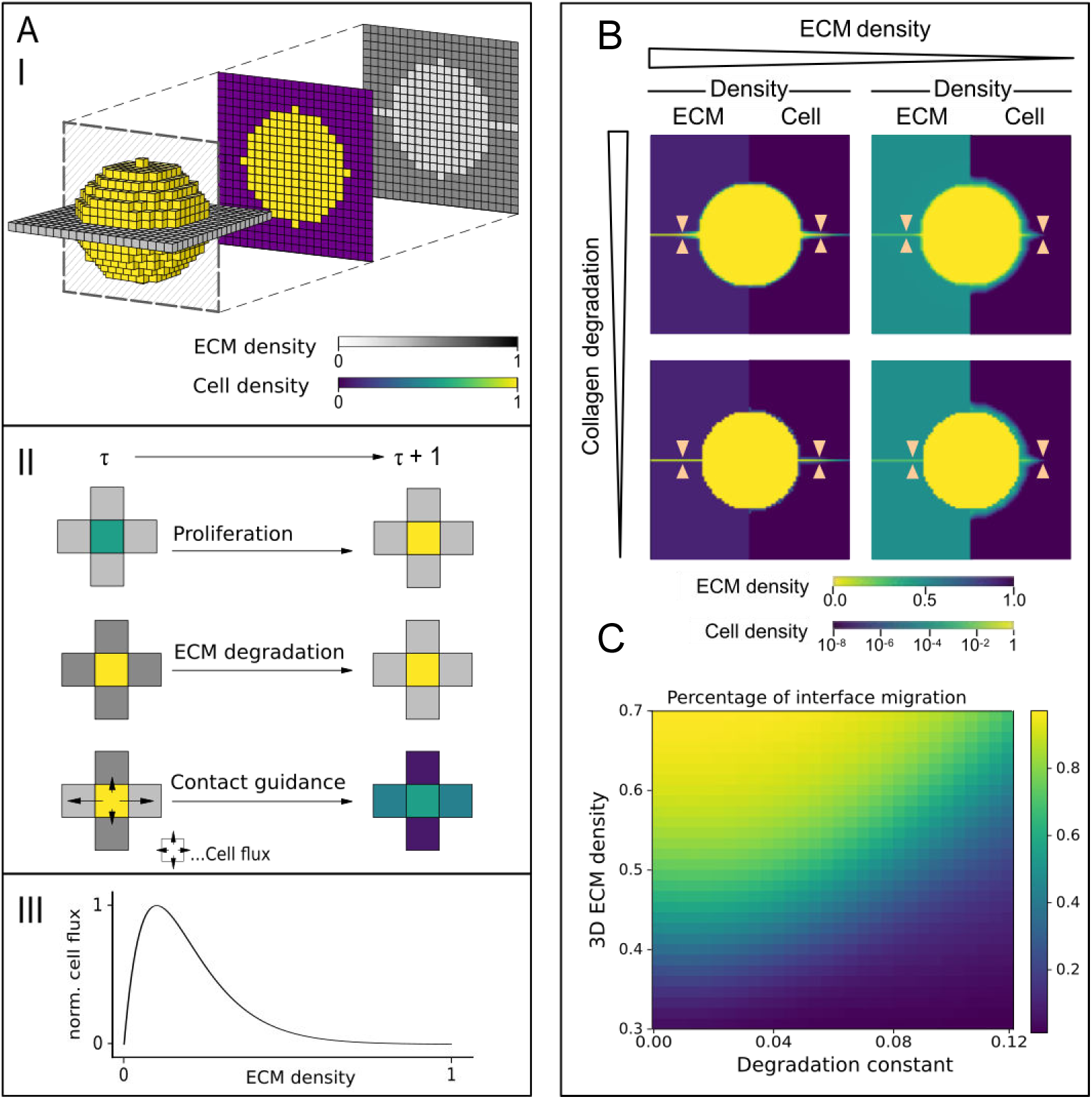
*In silico* cellular automata-based modeling of tumor cell invasion into a cleft. Development **(A)** and application **(B**,**C)** of an *in silico* model predicting tumor cell invasion along a low density interface. (**A)** (I) 3D lattice, each voxel containing scalar values for cell and ECM density ranging from 0 to 1. Initial conditions of high-density cell spheroid and low-density ECM interface. (II) During a discrete timestep t → t+1, the ECM and cell densities change deterministically due to proliferation: cell density increases with time; ECM degradation: cells degrade ECM therefore lowering the ECM density in the neighboring voxels; Contact guidance: Cells adjust their local movement direction along axis of high ECM density. This is implemented as a cell density increase caused by cell flux (indicated by the length of the arrows) depending on the ECM density (III) in the target voxel. **(B)** Depiction of ECM density (left) and cell density (right) after 100 time steps at varying collagen degradation and ECM density. Orange arrowheads indicate the location of the interface. **(C)** Quantification of percentage interface-guided migration of total migration as a function of ECM density and ECM degradation.

### Non-proteolytic migration is enabled by extracellular matrix guidance

To evaluate the prediction of the model on impaired infiltration of random ECM of high density, but not along the interface, when collagenolysis was inhibited, we applied the broad-spectrum MMP inhibitor GM6001 and monitored invasion in both the high-density and low-density interface assays. In control culture, MV3 cells invading along the interface between high-density matrices extensively degraded collagen, while MMP inhibition reduced collagen degradation along the interface by approximately 90% (Fig. 4A). After MMP inhibition, interface-guided migration was decreased for both low and high matrix density by approximately 50% (Figs. S4A-C), consistent with moderate hindrance imposed by the matrix [16]. The remaining invasion along the interface bordered by both high- and low-density matrix occurred predominantly as collective strands with subcellular protrusions of great length (Fig. S4A). Even with strongly diminished ability to degrade matrix, a 1.5-fold increased cell number and area covered along the cleft bordered by high-density compared to low-density matrix indicated an overall increased migration efficiency (Figs. 4B-D). By contrast, the maximum distance of the strand leader cells from the spheroid edge was marginally increased along the cleft bordered by low-density collagen compared to high-density matrix, indicating that the strong strand-like migration phenotype in the low-density assay enabled residual non-proteolytic migration (Figs. 4E; S4C). Non-guided migration into random matrix after MMP inhibition was abolished in high-density, but not in low-density collagen matrix (Figs. 4F-H; S4D,E). As end-point in high-density random collagen, cytoplasmic protrusions reached out of the core into the matrix whereas nuclei remained in the spheroid core (Fig. 4F, left column). Residual migration into low-density collagen was maintained by occasional strand-like cell patterns that also included cell nuclei (Figs. 4F, right column; S4D,E).

**Figure 4.**
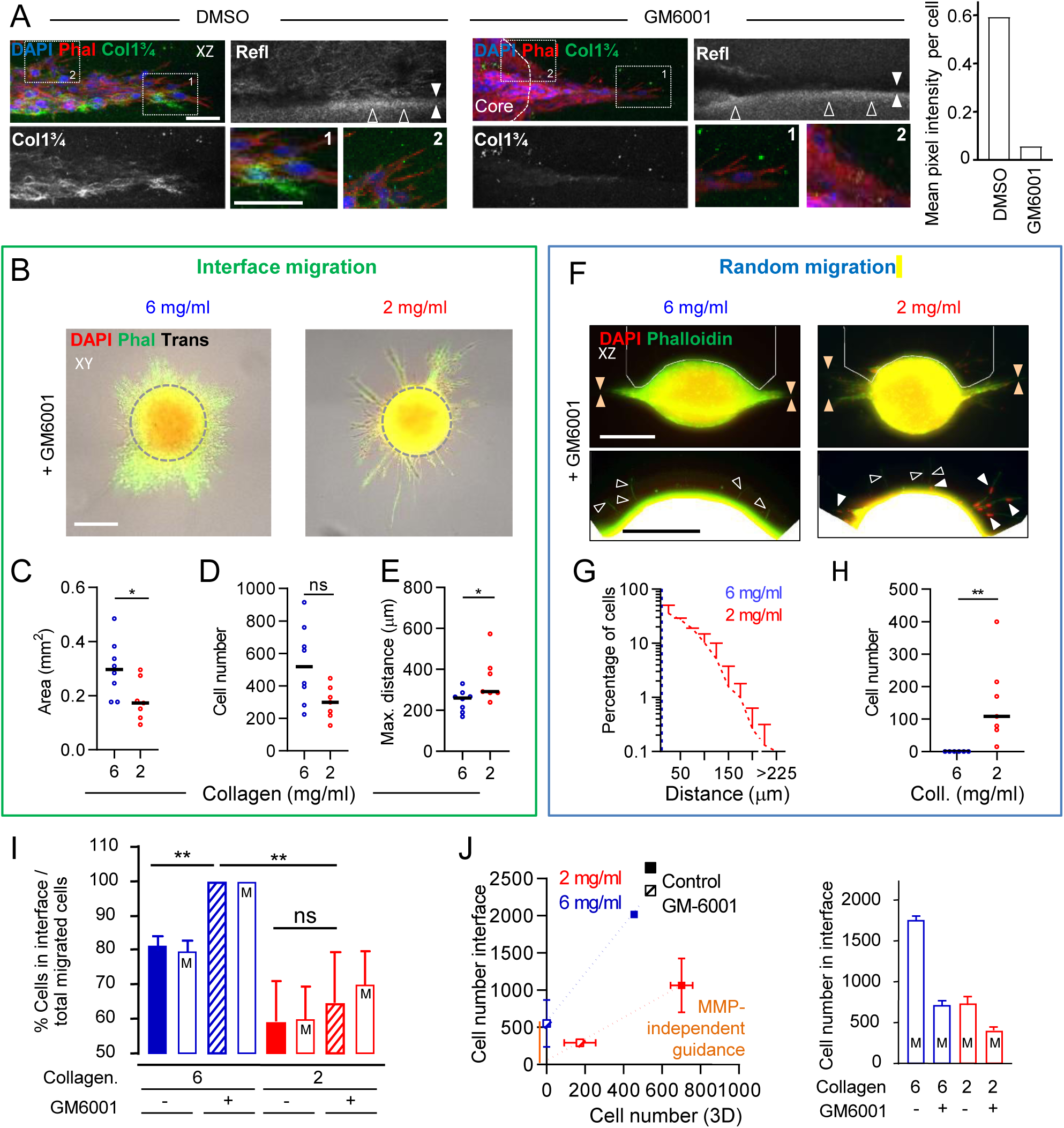
Differential MMP-dependence of invasion along interface or into random collagen. Cell emigration patterns in the presence or absence of MMP inhibitor GM6001 after 48h culture. **(A)** Xz view of interface migration at indicated conditions and in 6 mg/ml collagen after 48h culture in the presence or absence of GM6001. Where present, the rim of the core is indicated by white dotted line. Graph, COL1 ¾ signal per cell (counted as 75 and 118 nuclei per condition from the invasion zone, but not the core; n=1). **(B-G)** Cell emigration parameters in the presence of GM6001 after 48h. **(A)** Gray dotted circles indicate size of depicted spheroid when imaged at 0h culture. **(F)** Area bordered by white line in top images indicates the area of the zoom-in in the respective lower image; orange arrowheads, interface; open arrowheads, cytoplasmic extensions; closed arrowheads, invaded nuclei. **(C-E, G, H)** Quantification of interface-guided and non-guided migration parameters as described in Fig. 1 and Fig. 2G. Data represent 1-4 measurements (dots/line) per condition per experiment (N=2). **(I**,**J)** Interface-mediated versus non-guided migration in the absence or presence of MMP activity. **(I)** Percentage of emigrated cells in the interface compared to total number of emigrated cells per spheroid. **(J)** Total numbers of interface-guided cells plotted against non-guided cells. Orange line indicates condition of MMP-independent guidance. **(I, J)** Data from bars depicted with “M” were derived from *in silico* cellular automata-based model. *In vitro* data were derived from experiments depicted in Fig. S4B,E; data represent 1-4 measurements (bar/square) per condition per experiment (N=2). **(C-E, H)** Solid line, median, **(I**,**J)** Bars and squares, mean; whiskers, SD. Mann-Whitney test with, when necessary, Holm-Sidak, ns: not significant; *: P-value < 0.05; **: P-value < 0.01. Scale bars: **(A)** 50 μm; **(B, F)** 250 μm.

In summary, non-proteolytic migration into random matrix depended on matrix density in accordance to the physical limit of migration [16]. In contrast, guiding cues present within the high-density matrix, and to lower degree in the low-density assay where the guiding matrix cues were poorly defined, enabled persisting non-proteolytic migration. Hereby, the *in vitro* data confirmed the *in silico* prediction (Fig. 4I) that even without MMP activity, migration along the interface between high-density tissue would be less inhibited compared to low-density matrix, and even be most efficient (Fig. 4J). Consequently, the ratio of cells migrating along the cleft was unchanged upon MMP inhibition in the low-density assay, while emigration from the spheroid shifted completely towards interface-guided migration in the high-density assay (Fig. 4I).

## Discussion

We here developed an *in vitro* spheroid invasion assay consisting of two layers of 3D porous randomly organized collagen of high- or low-density, which formed a guiding track of confined 3D topology with lower collagen density. By a cross-section approach, cell invasion was effectively visualized, allowing to quantify the impact of substrate density, geometry and MMP-dependent collagenolysis on decision-making in invasion outcome for all ECM compartments. *In vitro* and *in silico* data show that the presence of a low-density area surrounded by restrictive matrix (1) supported cell motility and (2) enabled MMP-independent migration.

### Confining cleft of low ECM density facilitates migration

Our study demonstrates that migration occurred predominantly along clefts of low ECM density, especially when supported and bordered by high-density matrix. *In vivo*, tumor invasion is guided by cleft-like spaces with cross-sections ranging between 0.25 and 900 μm^2^ between larger structures such as blood vessels, nerves and thick fibers in interstitial tissues [4,6]. *In vitro*, we probed the lower range of this parameter space by modulating restrictive pore sizes in relation to the cross-section of migrating mesenchymal cells (around 50-100 μm^2^) [16]. Our results show an around 2-4 times increased pore size range within the cleft, with the high-density assay displaying larger differences in collagen density than the low-density assay, which consequently favored cell migration along the cleft.

In addition to the area of increased porosity, the cleft was bordered by a rim of very high density and low porosity, hereby confining migrating cells and increasing local ECM stiffness, especially in the better defined interface in the high-density matrix assay [14,16]. In both the high- and low-density spheroid invasion assay, collective migration along the interface was observed, either as a high cell density sheet or as thin collective strands harboring less cells, respectively, in accordance to previous observations of collective migration along a confined space [14,19]. Confinement supports and guides cell migration via cellular polarization and induction of the actin-myosin network alignment in the direction of the cue [25,28–30]. In addition, cells locomoting within stiffer 3D substrates increase integrin clustering, polarization of the cytoskeleton mechanotransduction, promoting migration rates in a pore size-independent manner [31–33]. These findings from the literature regarding migration-promoting biophysical properties of interface together with the predictions by the cellular automata model jointly support the strong cell guidance along the interface bordered by high-density compared to the low-density matrix.

### Role of collagenolysis in matrix invasion

Confined migration of mesenchymal cells within 3D dense matrices, combined with partial breakdown of surrounding matrix, creates as much space as necessary to accommodate invasive growth [14,15]. Matrix degradation and proliferation are coupled such that collagenolysis creates the necessary space for cell expansion [34]. This especially applies for cell migration in very dense matrices, where MMP activity is mandatory for the cell to move forward [16,34,35]. Interestingly, the density of the matrix determines the amount of collagenolysis to occur by a “digest-on-demand” mechanism, where higher matrix density results in more matrix degradation [27], and this rule was confirmed by the here developed *in silico* model. Cancer cells with active MMP-systems can thus invade a broad range of tissues, even within restrictive high-density areas. However, in the absence of MMPs, migration into dense 3D spaces is delayed or even abrogated [16]. Here, only cytoplasmic extensions, but no cell nuclei, reached the surrounding dense ECM, defining the non-guided migration into random dense matrix as MMP-dependent migration. By contrast, migration along the low-density clefts between high-density matrix persisted in the absence of collagenolysis, and enabled us to identify tissue clefts as an MMP-independent invasion niche. Matrix guidance by a low ECM density area within a dense extracellular matrix supports migration in both the presence of absence of MMP activity, thereby defining guidance as a primary determinant of cell patterning above matrix degradation.

### Implications

Locomotion into clefts *in vivo* may persist in the absence of collagenolysis and invasion into cleft-like matrix structures may thus not be responsive to protease inhibitor therapy, while invasion into densely structured randomly organized tissue is predicted to be impaired upon MMP inhibitor treatment. Our data thus imply that realistic 3D ECM models probing cancer invasion should match *in vivo* tissues with regard to biophysical properties including varying density and complex geometry. The spheroid invasion assay mimics cleft-like low-density tissue conduits filled with interstitial fluid, small vessels or low-density networks of fibrillar collagen, glucosaminoglycans, or other ECM components providing spaces of reduced resistance in interstitial tissues [6]. Therefore, the interface assay can be applied, and further developed, to dissect the impact of complex tissue heterogeneities *in vivo* on cancer cell invasion, thereby supporting and reducing animal experiments.

---

**Supplementary Figure 1.**
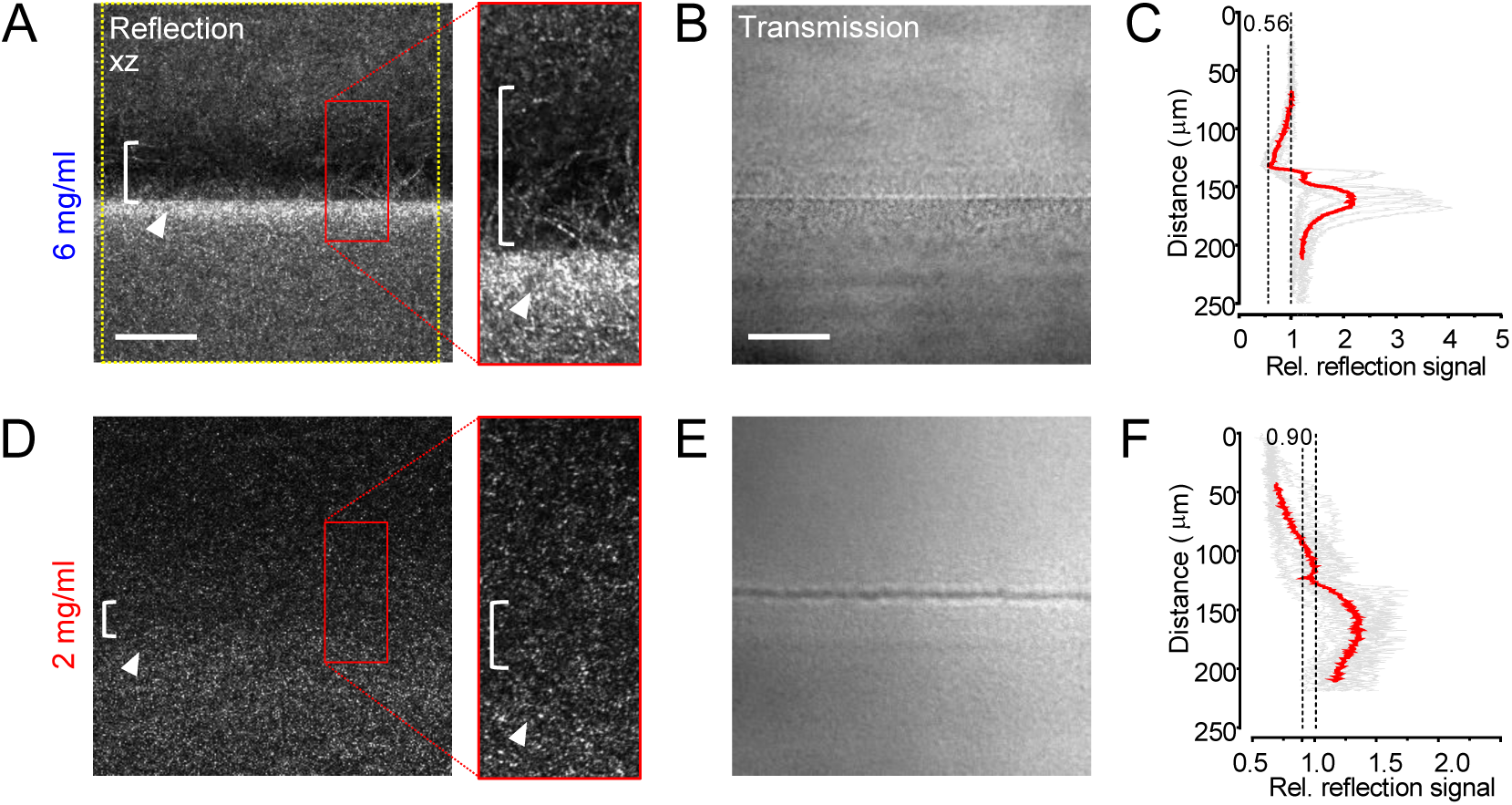
Characterization of the collagen-collagen interface. Reflection-based analysis of collagen signal of manually cross-sectioned lattices. **(A**,**D)** Collagen-collagen interface between collagen lattices of indicated concentrations. Arrowheads indicate regions with highest collagen density; brace indicates area with decreased collagen density. Collagen reflection signal, side view. Yellow square indicates the 1500 pixel-wide area used for quantification, and red rectangles the regions of the insets shown on the right. **(B, E)** Corresponding transmission signals. **(C, F)** Calculation of cleft versus random matrix density ratios. Quantification of the mean gray value along a 1500 pixel wide line from top to bottom of image perpendicular to the interface (indicated in A by yellow square) aligned along the interface. N=3; 4-5 images per experiment. Red line, mean; gray lines, individual experiments. Scale bar: 50 μm; inset: 10 μm.

**Supplementary Figure 2.**
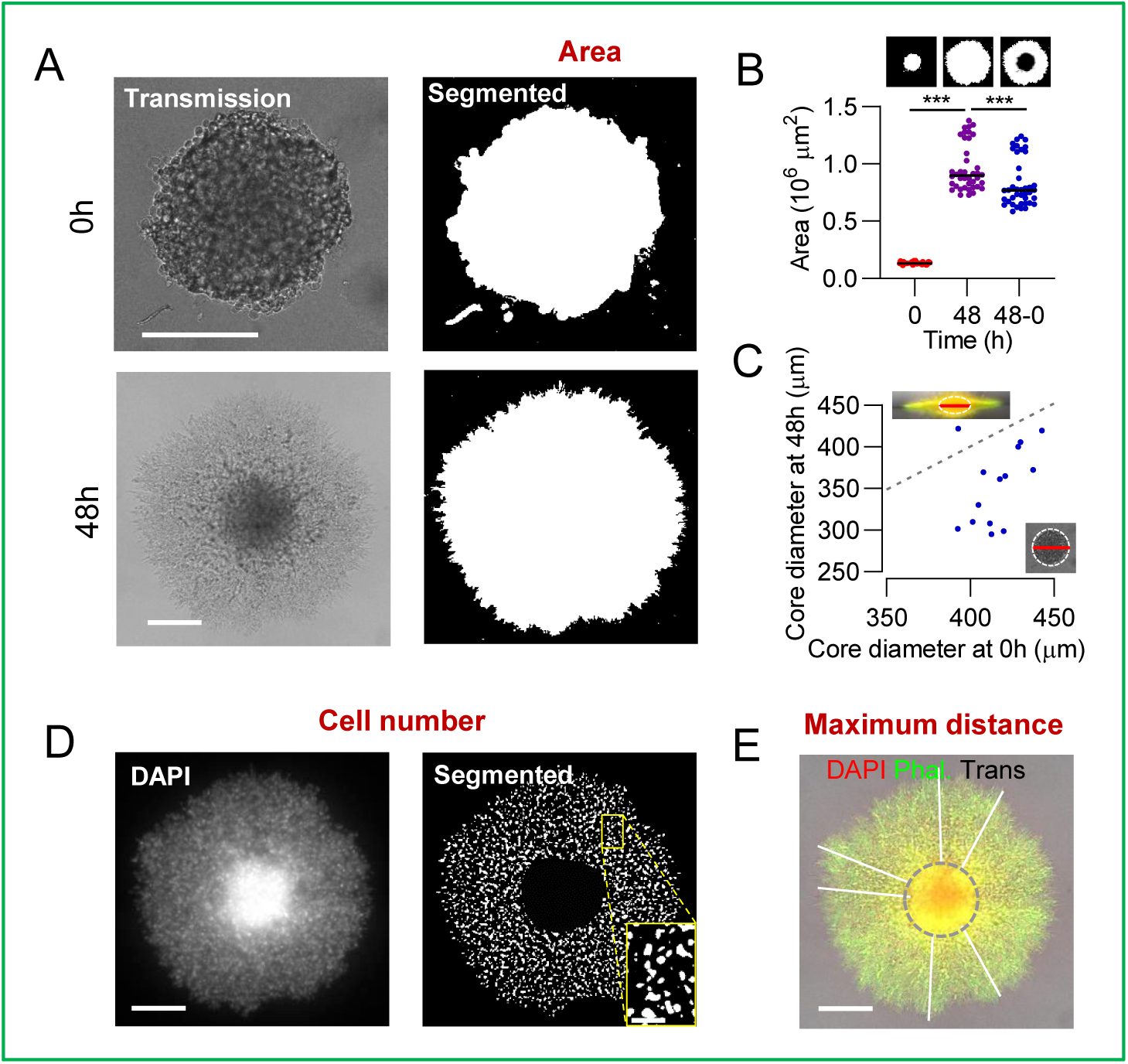
Strategies to quantify migration parameters for interface migration. Approaches for the quantification of the parameters ‘Area’, ‘Cell number’ and ‘Maximum distance’ of interface-guided migration. **(A)** (Left) Original spheroids in collagen and (right) automated segmentation of the cell-populated area at 0h and at 48h culture. **(B)** Quantification of segmented indicated areas. **(C)** Spheroid core areas after 0 and 48h culture were measured and compared. Shrinkage of the spheroid core area after 48h culture was detected (see gray diagonal line which indicates y=x; as well as orange circle in Fig. 1K for visualization). **(D)** Automated segmentation of DAPI signal after manual exclusion of the core. Left, original image; right, corresponding segmentation. **(E)** Depiction of manual distance quantification from the rim of the spheroid to the six furthest migrated cells. **(B, C)** Each dot represents a spheroid; median, solid line. Mann-Whitney test with followed by Holm-Sidak, ***: P-value < 0.001. Scale bars: **(A, D-E)** 250 μm, **(D inset)** 50 μm.

**Supplementary Figure 3.**
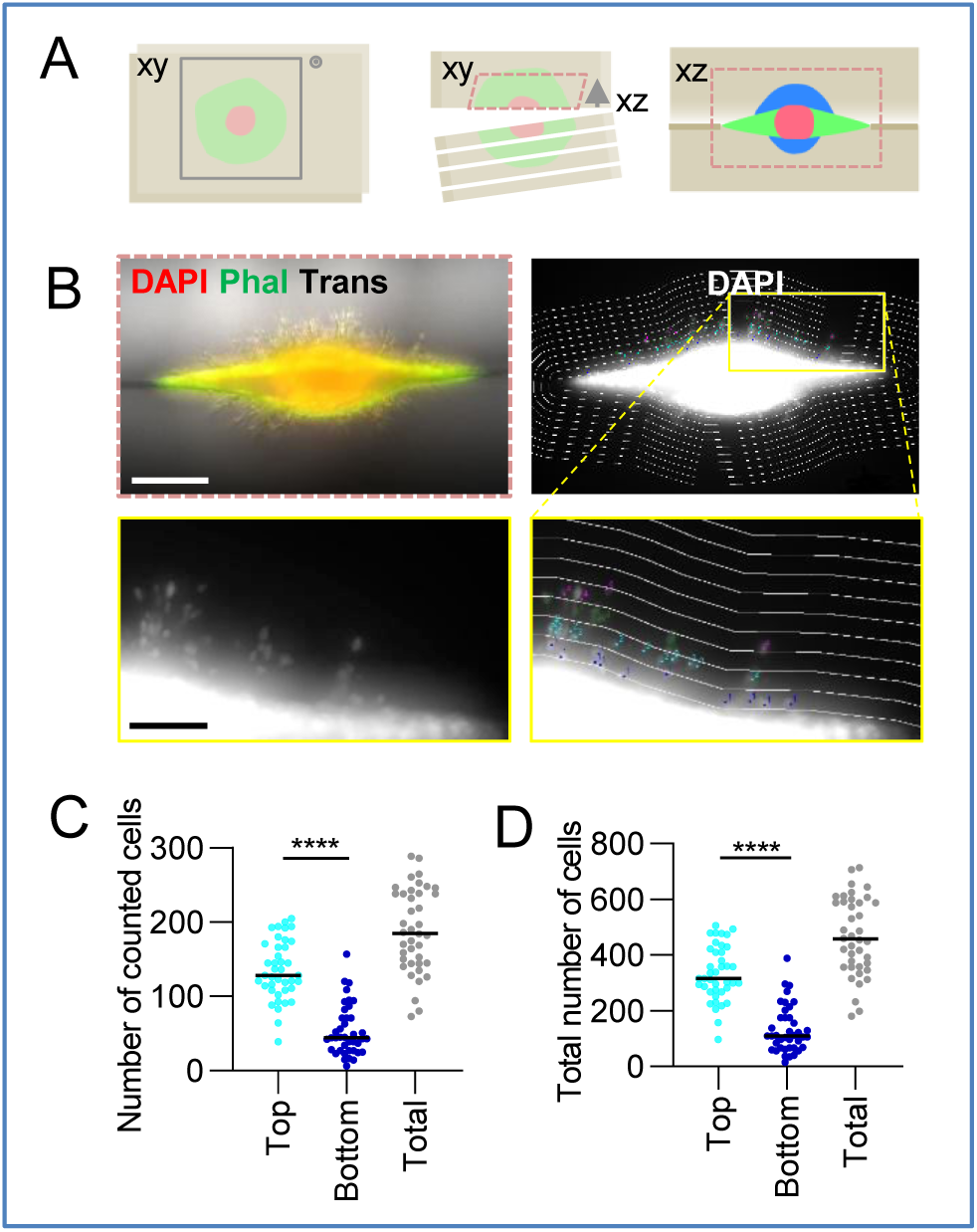
Strategies to image and quantify non-guided migration. 48h cell cultures were manually dissected to visualize and quantify cells that migrated into the random collagen beyond the interface. **(A)** Cartoons depicting imaging strategy before and after manual cross-sectioning. Not all cells within the 3D matrix were imaged as only one cross-sectioned lattice per complete lattice, and thereby half of the spheroid, was imaged, in addition to the limited penetration depth of the used imaging method (red dotted rectangle in the middle cartoon depicts imaged area). Using the penetration depth, non-guided migration depth and core size, it was calculated that only 40.4% of non-guided cells were imaged. Therefore, the total number of non-guided cells was corrected with a factor 2.47. **(B)** Cells migrating into random collagen (6 mg/ml) beyond the interface after imaged after manual cross-sectioning of the fixed cell-collagen 48h culture. Right, binning of random collagen regions into 25 μm sections created by expanding ROI’s based on manual count of widefield nuclear fluorescence signal. Images are overlays of 24 z-stacks of each 10 μm distance, resulting in 230 μm deep widefield images. A cell was counted per bin when at least half of the cell nucleus was located in the area. White lines indicate edges of sections; colors represent cells in different distance sections. Bottom images are zoom-ins. **(C)** Number of counted cells per 230 μm deep top and bottom layers. **(D)** Total number of cells per layer after correction with a factor 2.47 for non-imaged, and therefore non-counted, cells. From here on, all cells within the random collagen compartment were multiplied with this correction factor. **(C, D)** Each dot represents a spheroid, median, solid line. Mann-Whitney test, ****: P-value < 0.0001. Scale bar: 250 μm, inset: 100 μm.

**Supplementary Figure 4.**
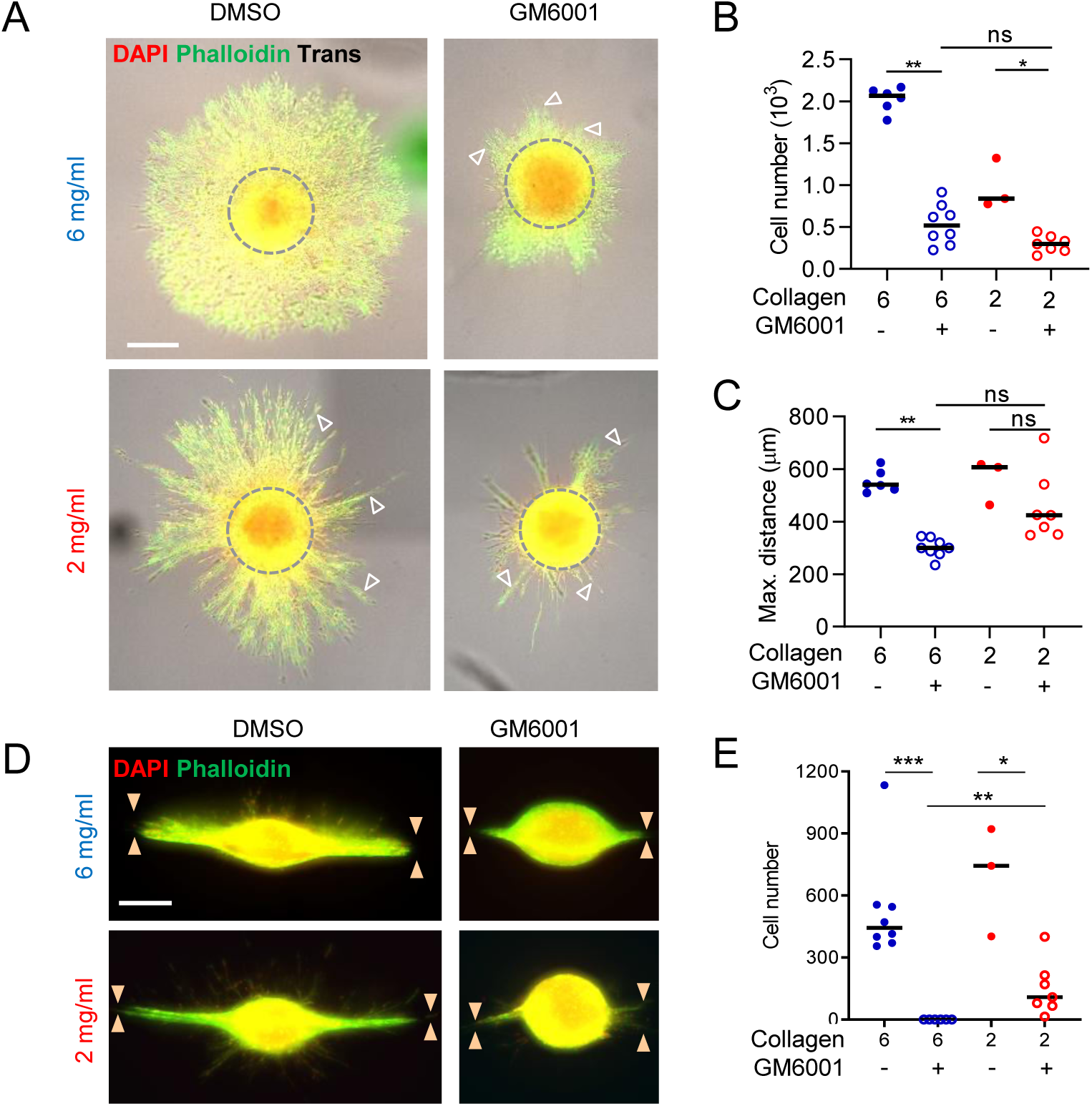
The ratio of interface-guided migration and invasion into random collagen depends on MMP-dependent collagen degradation. Images and data of the GM6001 condition from Figs. 4B, D, E, F and H and are here paired with the respective controls. **(A**,**D)** Depiction and **(B**,**C**,**E)** quantification of cell emigration into the interface **(A-C)** or fibrillar collagen **(D**,**E)** in the absence or the presence of MMP inhibitor GM6001. Arrowheads indicate strand-like migration **(A). (B**,**C**,**E)** Data represent 1-4 measurements (dots) per condition per experiment (N=2). Each dot represents a spheroid, median, solid line. Mann-Whitney test followed by Holm-Sidak, ns, not significant; *: P-value < 0.05; **: P-value < 0.01. Scale bars: 250 μm.

